# Conflict-Mediated Group Size Regulation: A Theory of Supraoptimal and Suboptimal Group Size

**DOI:** 10.64898/2026.06.26.734857

**Authors:** Eric Schniter

## Abstract

Observed group sizes rarely match the size that would maximize what each member gets from belonging. We propose a two-part theory in which group size is regulated by two related conflicts: insider-outsider conflict over admission, and within-group conflict as crowding, competition, and social tensions intensify with size. Three strategies are available within a group of interchangeable members: admission, exclusion, and fission. The models hold each member’s standing and resource exploitation strategy constant. While both vary and determine a member’s own game and its payoffs, explaining this complexity and its consequences is beyond our scope. The theory’s first part shows that even when exclusion is unavailable, fission dynamics alone drive group size away from the optimum in both directions, with the pattern set by how prospective joiners encounter groups and by the geometry of fission. When joiners compare groups across a shared landscape and fission is asymmetric, the standing distribution is bimodal: supra-optimal large groups coexisting with a sub-optimal mode of small groups, the pattern characteristic of fission-fusion societies. The second part promotes exclusion and fission to active decisions: incumbents weigh the per-capita cost of accommodating entry (*β*) against the costs of coordinated exclusion (*c* + *γN*\*) and fissioning (*F*). Two inequalities of the same form do the work, each setting a payoff slope against the slope of a coordination cost. *β* > *c* + *γN*\* partitions populations into two regimes: where it holds, exclusion is viable and groups lock at the optimum size; where it fails, groups grow past the optimum and cycle through recurrent fission. *α* > *φ* then fixes the geometry of that fission, separating splits in which members leave independently, which strand small daughters and give the bimodal distribution, from splits in which several members depart at once. Modal group size, fission frequency, and exclusion behavior together identify which regime governs a population — a set of predictions applicable across fishes, social insects, birds, and mammals including primates and human foragers.

## 1. Introduction

Understanding why animal groups deviate from their optimal size is one of the enduring problems in behavioral ecology. The benefits of group living, such as shared vigilance, cooperative foraging, and collective defense, are well established, as is the observation that per-capita returns from these benefits follow a hump-shaped function of group size: rising initially as groups grow, then declining as crowding and within-group competition erode the gains. The group size at which per-capita returns are maximized is the nominal optimum, *N\**. Yet observed group sizes routinely deviate from *N\**, in both directions and often substantially, and the conditions under which they do so remain incompletely understood.

The supraoptimal case (groups larger than *N\**) is the more theorized of the two. Sibly (1983) demonstrated that *N\** is individually unstable: groups grow beyond it through individually rational joining and settle at a larger Nash equilibrium size. Sibly’s result has become a cornerstone of group size theory and is broadly supported empirically.

Sibly’s instability result was challenged shortly after publication by Giraldeau and Gillis (1985), who showed that whether *N\** is stable depends on the shape of the per-capita fitness function: when the declining slope is steeper than the rising slope, groups do not grow past the optimum. We make this qualitative insight quantitatively precise below. Kramer (1985) extended the same debate to colonial breeders, a paradigmatic empirical test case for the supraoptimal-group hypothesis.

The suboptimal case, groups smaller than N*, is empirically well documented and has been theorized mainly through exclusion. In wild baboon troops, for example, small groups bear greater costs relative to intermediate sizes (Markham et al., 2015). Higashi and Yamamura (1993) formalized insider–outsider conflict as a strategic membership game in which equilibrium group size depends on the costs of joining versus exclusion: groups may settle below N* when the cost of defending the optimum exceeds the cost of preserving a smaller group.

We argue that this framing overlooks a mechanism capable of generating suboptimal modal group size without any exclusion at all: the interaction between fission dynamics and joiner pool ecology. When a group splits, it produces a smaller resulting daughter below *N*\*; the larger may remain above it, particularly when the split sheds only a small number of members. What happens next depends critically on whether prospective joiners arrive independently at each group from local catchments, or whether they choose among all available groups across a shared landscape. Under local joiner pools, daughters grow autonomously and modal group size settles above *N*\* — consistent with Sibly. Under a global joiner pool, daughters compete for the same joiner stream; if the split was symmetric this drives both daughters toward the fission threshold, leaving modal size supraoptimal rather than sub. Sub-optimal sizes instead emerge when fission is asymmetric: a crowded group sheds a few members while the bulk persists, and the resulting small daughter, far below *N*\*, is the least attractive group in the pool and is bypassed by joiners while large groups continue to recruit. Suboptimal modal group size emerges from fission dynamics alone, with no exclusion required, when fission is asymmetric rather than symmetric.

We develop the framework in two stages. The first treats fission as the sole regulatory mechanism and shows how the structure of the joiner pool (local versus global, preference-based versus random) determines whether the standing distribution of group sizes is unimodal near the optimum, bimodal with coexisting sub- and supra-optimal modes, or broadly over-dispersed. The same payoff function, fission threshold, and individual rationality assumptions produce opposite directional predictions depending on a single ecological parameter. The second extends the fission-only model by introducing exclusion as a second regulatory mechanism, derives a viability condition that partitions a parameter space defined by exclusion costs, coordination burden, and fission costs into two qualitatively distinct dynamic regimes, and shows how joiner pool ecology from the first part shapes which regime governs any given population. Together, the two models provide a unified framework for understanding the full range of observed group size deviations from the nominal optimum. Throughout, the group is a set of interchangeable members whose responses to size are to admit, exclude, or split. The models hold each member’s standing and resource exploitation strategy constant. In reality, both vary and determine a member’s own game and its payoffs, resetting the costs admission, exclusion, and fission operate on rather than acting within them; where this operates it can permit larger stable groups. Explaining this complexity and its consequences is beyond the present scope. Among non-human animals, resource availability and clustering interact with social standing (determined by dominance rank and association bonds) to shape each member’s exploitation strategy and its payoffs. Among humans, the same logic operates through resource exploitation strategy, and through leveling mechanisms and norms that affect a member’s standing, while kinship and coalition structure separately shape who plays the game with whom.

## 2. Background

### 2.1 Optimal group size and the payoff function

The theoretical foundation for group size research rests on a simple observation: per-capita benefits of group living are a non-monotonic function of group size. Pulliam and Caraco (1984) synthesized the early formal treatments, showing that per-capita returns rise from a solo baseline *R*_1_ as group size increases (e.g., because of shared vigilance, cooperative foraging, and collective defense) reach a maximum at the nominal optimum *N\**, and decline thereafter as crowding, within-group competition, and resource dilution intensify. The nominal optimum is thus the group size that maximizes individual fitness under steady ecological conditions.

Throughout this manuscript, the per-capita payoff function *P*ᵢ(*n*) is treated as a component of inclusive fitness attributable to group membership at size *n*, following the phenotypic gambit (Grafen, 1984). Empirical tests of this framework span taxa from fish shoals (Buston, 2003; Pitcher, 1986) to social insects (Grinsted and Field, 2018a) to cooperatively breeding birds (Sturrock et al., 2022) to rodents (Waterman, 2002) to primates (Markham and Gesquiere, 2017) to human foragers (Smith, 1981). The general shape of the payoff function is well supported, though estimates of *N\** vary substantially across populations and ecological contexts (Pulliam and Caraco, 1984; Rodman, 1981). A persistent challenge is explaining the systematic bidirectional divergence of realized modal group size from *N*. Recent reviews of the primate literature have flagged this divergence as an unresolved puzzle: long-term monitoring of wild populations shows groups persisting above their predicted optima, despite the theoretical expectation that prospective joiners should preferentially affiliate with optimally-sized groups (Markham and Gesquiere, 2017). Among human foragers, parallel puzzles have motivated active inquiry: the average forager group size far exceeds any non-human primate analog (David-Barrett, 2023). Here we develop a theoretical account showing that persistence above and below the optimum follows from joiner pool ecology and the costs of coordinated exclusion.

### 2.2 Supraoptimal group size: Sibly’s result and its extensions

Clark and Mangel (1984) elaborated this logic in a more general setting, and the result has since been treated as the canonical prediction for social groups in which members cannot or do not exclude newcomers. It is broadly consistent with empirical findings that realized group sizes frequently exceed the group size at which per-capita fitness is maximized (Smith 1981; Markham et al. 2015). This logic is especially relevant to fission-fusion species: groups that repeatedly split and regrow will spend most of their cycle above *N\**, approaching the fission threshold before splitting again.

### 2.3 Suboptimal group size: insider-outsider conflict and exclusion

Subsequent theoretical work has refined the conditions under which exclusion succeeds (Shen et al., 2023, 2017). Empirically, viable exclusion has been documented in taxonomically distant systems. A particularly clear case is the Cape ground squirrel (Waterman, 2002), in which matrilineal kin groups maintain a stable cap of approximately three breeding females through coordinated reproductive suppression of younger females, with fission occurring only when this active regulatory mechanism is overwhelmed by group growth — an empirical instance of viable exclusion holding group size below the size at which fission would otherwise be triggered. A second clear case is the clown anemonefish (Buston, 2003), in which dominant breeders regulate group size through forcible eviction of existing subordinates and active prevention of new recruitment, demonstrating viable exclusion in a phylogenetically distant taxon and through a different behavioral mechanism. But exclusion is costly, imperfect, and subject to free-riding among incumbents — conditions that limit how reliably it can hold group size at or below *N\**.

What the insider-outsider conflict framework does not address is the possibility that suboptimal modal group size could arise without exclusion at all, as a consequence of fission dynamics interacting with joiner pool ecology. This gap is one of the motivations for the present paper; we take it up alongside several additional problems that have remained unresolved in the joint analysis of admission, exclusion, and fission.

### 2.4 Fission as a regulatory mechanism: what the literature has and has not addressed

Fission, the splitting of an oversized group into two daughter groups, is recognized as the primary mechanism by which groups regulate upper size bounds in many taxa (Aureli et al., 2008b). Seno (2006) modeled group size dynamics via fission and fusion with inclusive fitness considerations, recovering several of the qualitative predictions of Sibly’s framework. The broader fission-fusion literature has examined the behavioral and ecological triggers of splitting events and the consequences of fission for social structure (Sandel et al. 2026; Feder et al. 2025).

A particularly direct theoretical precursor to the present framework is Smith’s (1985) analysis of Inuit foraging groups, which formalized the conflict of interest between current group members and prospective joiners. Smith showed that where the alternative to group membership is low-return solitary foraging, joiners prefer group sizes that exceed the per-capita optimum for incumbent members, with the conflict resolved in favor of joiner preferences in the absence of effective exclusion. We build on this foundation in two directions: by treating fission as an additional regulatory mechanism that responds to dilution beyond the joiner-admission decision, and by promoting exclusion from a possibility weighed against admission to an active decision subject to coordination costs and free-riding. We also formalize how joiner pool structure (i.e., whether prospective joiners can compare available groups or encounter them sequentially) interacts with the conflict to determine the magnitude and direction of deviation from the optimum. Smith’s relatedness weighting of payoffs is a natural extension within the inclusive-fitness framing adopted here: when *r* > 0, incumbents partially internalize fitness benefits flowing to related group members and related prospective joiners, raising the effective payoff of admission and attenuating the insider-outsider conflict that drives the present model’s dynamics. The baseline analyses below treat *r* = 0 to isolate the structural results, but the directional prediction is clear — higher within-group relatedness pushes modal group size upward relative to the *r* = 0 baseline.

A separate body of theory addresses group-size distributions from the opposite direction, specifying rates of splitting and merging directly at the population level rather than deriving them from individual decisions. Lerch and Abbott (2024) and the coagulation-fragmentation tradition (Ma et al., 2011) are complementary in scope here, both characterizing population-level group-size distributions from specified between-group rates. Safran et al. (2007) identified this individual/population separation as a major gap in the group-living literature and called for models that link the two; the present framework supplies that link, using individual admit-exclude-fission decisions to predict the population-level signatures these distributional approaches characterize.

Extending this individual-to-population link, an unresolved issue is how population-level group-size distributions emerge when fission interacts with different joiner ecologies. Earlier theoretical work, including Sibly’s original analysis, implicitly assumed that each group attracts joiners from a local catchment independently of other groups. Under this Case 1 assumption (which we examine first below before relaxing), post-fission daughters each grow autonomously and the population-level modal size reflects the fission-regrowth cycle of individual groups: above *N*\* for most of the cycle, settling near the fission threshold. This is the ecologically appropriate model for relatively isolated groups or for species in which prospective joiners encounter single groups rather than assessing multiple options simultaneously; Cases 2 and 3 below relax that assumption.

A parallel literature in archaeology and human ecology describes the same regulation in human settlements. Scalar-stress theory holds that within-group conflict rises with group size (Johnson, 1982), the declining slope captured here by β, and studies of early villages document recurrent fission as the mechanism that resolves this conflict until higher-level integrative institutions emerge, after which fission ceases and settlements grow large and stable (Bandy, 2004). The present framework formalizes what these accounts describe qualitatively, giving the criterion that separates the regime in which recurrent fission does the regulating from the one in which exclusion and integrative institutions hold a group together, and adds the joiner-pool distribution predictions and the post-fission divergence prediction they do not make.

## 3. The fission-only model

### 3.1 Setup

The fission-only model of group size regulation allows us to ask what determines modal group size when fission is the only available regulatory mechanism (i.e., exclusion is not viable). The declining slope of the payoff function, captured by the parameter *β* introduced below, already embeds the ambient costs of group life (e.g., within-group competition, conflict, crowding, and disease) as passive costs that accumulate automatically with group size. The fission-and-exclusion model builds on this fission-only foundation by asking what happens when groups can actively respond to those costs through exclusion, treating membership regulation as a strategic collective-action problem rather than a fixed environmental parameter, while fission remains governed by the threshold and cost parameters established in Part 1.

### 3.2 The payoff function

Let *R*_1_ denote the per-capita return available to a solitary individual — the outside option against which group membership is evaluated. Let *P*_I_(*n*) denote the per-capita payoff of an intact group of size *n*. We model *P*_I_(*n*) as a piecewise linear function in equation (1), consisting of two line segments meeting at the nominal optimum *N\**: a rising segment below *N\** with slope *α*, and a declining segment above *N\** with slope *β*:

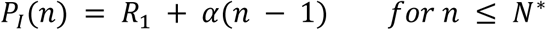

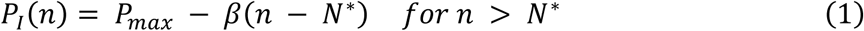

where *P*_max_ = *R*_1_ + *α*(*N\** − 1) is the peak per-capita payoff at the nominal optimum. The two slopes *α* and *β* are independent positive parameters. When *α* = *β* the function is symmetric, we get the classic tent shape used in most prior treatments. When *α* ≠ *β* the function is asymmetric: the rate at which membership becomes more valuable as the group grows toward *N\** differs from the rate at which it becomes less valuable as the group grows beyond it. Both slopes are empirically meaningful and, as we show below, their ratio determines the direction of modal group size deviation from *N\** under local joiner pool conditions. The piecewise linear form is adopted for analytical tractability; qualitatively identical results hold for smooth, continuously differentiable payoff functions with the same slope asymmetry.

The two slopes capture a distinction that the symmetric model conceals: the processes that make groups valuable as they grow need not mirror the processes that make them costly as they grow further. On the rising side, benefits such as shared vigilance, cooperative foraging, and collective defense accumulate as new members contribute to public goods. On the declining side, costs such as within-group competition, conflict, crowding, and disease transmission accumulate as the group exceeds the size at which resources and social tolerance are comfortably shared. There is no theoretical reason these should be symmetric, and in practice they rarely are.

### 3.3 Sources of payoff asymmetry: why α and β differ

Understanding what drives slope asymmetry matters because, as the fission threshold derivation below shows, the ratio *α*/*β* determines the direction of modal group size deviation in a local joiner pool system. Giraldeau and Gillis (1985) first noted that slope asymmetry governs the direction of deviation from *N*\*. We make their qualitative insight quantitatively precise by deriving the analytical relationship between *α*, *β*, and modal group size. Two broad classes of asymmetry generate the two directions of deviation; each is illustrated by familiar empirical systems.

#### 3.3.1 Steep decline, shallow rise (β > α): costs escalate faster than benefits accumulate

Several well-documented mechanisms generate the *β* > *α* asymmetry: density-dependent pathogen and parasite transmission, whose contact rates scale roughly with the square of group size (Altizer et al., 2003); intensifying within-group competition for rank and resources, with measurable fitness costs (slower infant development, longer interbirth intervals) in larger primate groups (Borries et al., 2008); and range-mediated predation exposure, as in Cape ground squirrels, where juveniles foraging farther from the refuge burrow suffer disproportionate mortality (Waterman, 2002). In each, the payoff function falls more steeply than it rises and the fission threshold is reached sooner than the symmetric model predicts.

#### 3.3.2 Shallow decline, steep rise (α > β): benefits accumulate faster than costs escalate

The opposite asymmetry, *α* > *β,* arises where early members add large marginal benefits that saturate quickly while crowding costs stay flat across a wide supraoptimal range — as in cooperative predator defense, where each additional defender matters most at small group sizes (McNamara and Houston, 1992), and cooperative breeding, where scarce helpers contribute disproportionately to reproduction (Grinsted and Field 2018; Sturrock et al. 2022). This raises the fission threshold above the symmetric baseline, so groups grow larger before splitting.

### 3.4 How α/β maps onto conflict

The *α* and *β* parameters have a natural interpretation in terms of two related conflicts that arise as groups grow. Below *N*\*, adding a member benefits both incumbents and the newcomer at rate α, such that interests are aligned and there is no conflict. At *N*\* and above, two conflicts emerge simultaneously and are both captured by *β*. The first is an insider-outsider conflict: adding a member benefits the joiner (whose alternative is solitary low-return existence) but harms each incumbent by *β* per added entry. The second is a within-group conflict: as the group exceeds *N*\*, all current members face accumulating costs of crowding, competition for resources, social tension, and disease transmission, growing at the same rate *β*. The two conflicts are coupled, each new joiner intensifies within-group conflict for everyone present; but they are conceptually distinct, and the three mechanisms developed in this paper target them differently.

The two models in this paper formalize three responses, distinguished by whether incumbents passively accept the resulting costs or actively bear a coordination cost to resolve them. Admission is a passive response to insider-outsider conflict: when *n* < *N\**, it realizes mutual benefit, and when *n* ≥ *N*\*, it accommodates the joiner’s preference and lets each incumbent absorb the *β* cost rather than resolve it. Exclusion is the active counterpart: incumbents pay the per-capita coordination cost *c* + *γN*\* to suppress insider-outsider conflict by enforcing their preference against further entry. Fission primarily targets within-group conflict, dissolving the shared social space when accumulated within-group costs exceed the per-capita cost of splitting; per-capita payoff recovers in each daughter, and within-group conflict is distributed between the now-smaller groups. The viable exclusion condition, given by equation (2), identifies when exclusion is preferred over passive accommodation; bringing the cost of fission into that comparison then determines when fission is preferable to either response.

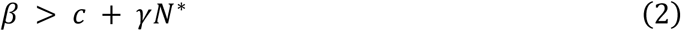

### 3.5 The joiner condition and why exclusion fails

A prospective newcomer joins if and only if group membership yields a higher return than the outside option. This gives the joiner entry condition:

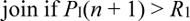

On the rising side of the payoff function this condition is always satisfied for *n* < *N\**. On the declining side it remains satisfied until *P*_I_(*n*) returns to *R*_1_, which occurs at the Nash equilibrium size given in equation (3), one entrant below the size at which an incumbent’s payoff would itself fall to *R*_1_ (a joiner evaluates the post-entry payoff *P*_I_(*n* + 1), not *P*_I_(*n*)).

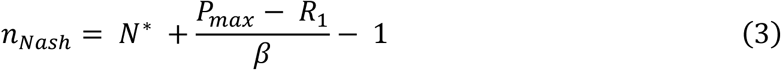

In the symmetric case this is 2*N*\* − 2; with a steeper decline (*β* > *α*) it is closer to *N\**; with a shallower decline (*β* < *α*) it is further above *N\**. Joining remains individually rational across this entire range. The admit option in the fission-and-exclusion model below adopts the same convention: the payoff to admitting one more member at size *n* is *W*_A_(*n*) = *P*_I_(*n* + 1), so the viable-exclusion comparison there is already evaluated post-entry.

Incumbents prefer the group not grow beyond *N\**, since per-capita payoff declines above the nominal optimum. But any single incumbent considering resistance faces an asymmetric cost structure: exclusion costs (e.g., monitoring, signaling, aggression, coalition-building) fall privately on the excluder while the benefit of a less diluted group is shared by all. Exclusion is therefore a public good, undersupplied by individually rational actors. Without a mechanism for coordinating joint exclusion, the group cannot resist entry. The fission-and-exclusion model addresses what happens when such coordination becomes possible; in the fission-only model it does not exist.

### 3.6 The fission threshold

The fission threshold, given by equation (4), follows directly from solving *P*_I_(*n*/2) > *P*_I_(*n*) for the piecewise linear payoff function; the full derivation is given in Appendix A.

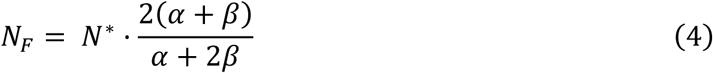

The model treats fission as a symmetric split into two comparable daughters of roughly half the parent’s size. This is a tractable idealization at one end of a continuum; at the other end a single individual dissociates, sometimes drawing followers, to form a much smaller group that may later re-fuse. The threshold logic does not require the split to be exactly equal — asymmetric or partial fission alters the post-split size distribution and how quickly the regrowth cycle resets, but not the existence of a size above which splitting becomes individually rational.

### 3.7 Departure geometry and the second viability condition

The threshold above fixes when a group splits but not how its mass divides. Let *m* denote the number of members who leave together, 1 ≤ *m* < *n*, and let the departing set bear a coordination cost of the same form as the exclusion cost introduced in the second model:

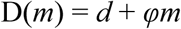

with *d* the baseline cost of organizing a departure at all and *φ* the rate at which that cost rises with the number who must be held to a joint decision to go. The micro-foundation is the one that governs exclusion (Appendix C): a volunteer’s dilemma whose probability of successful coordination falls as the number of required contributors rises.

A member who leaves in a set of size *m* receives the payoff of the group that set forms, net of the cost of assembling it and of fission itself:

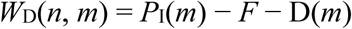

Departing sets form at or below the optimum, so *P*_I_(*m*) = *R*_1_ + *α*(*m* − 1), and the payoff to a departing member is *R*_1_ + *α*(*m* − 1) − *F* − *d* − *φm*. Its derivative with respect to *m* is *α* − *φ*, and that single sign fixes the geometry of every split the model produces. Where *φ* > *α*, each member added to a departing set costs more to bring along than the rising limb of the payoff function returns: the payoff to leaving falls with *m*, and departure goes at its smallest unit, one member leaving alone if need be. Where *φ* < *α*, each added member returns more than it costs, the payoff to leaving rises with *m*, no member departs alone, and departure waits until a set assembles. The set then grows until further recruits would dilute it rather than strengthen it, which is at *N*\*.

*α* > *φ* is therefore a second viability condition, standing to the geometry of fission as *β* > *c* + *γN*\* stands to the choice between admission and exclusion. Both compare a slope of the payoff function against the slope of a coordination cost, and in both it is the payoff slope winning that makes coordinated action viable. The derivation is given in Appendix E.

Two consequences follow, and both concern results stated elsewhere in this paper. The symmetric halving retained for *N*_F_, *N*_AF_, and *n*_EF_ is the *φ* < *α* branch at the size where the two coincide: a departing set of *N*\* leaving a parent of 2*N*\* leaves *N*\* behind, so the split is even. Away from that size the same branch predicts an uneven division, with the departing daughter at *N*\* and the remainder at *n* − *N*\*. The asymmetric fission of Case 2 below is the *φ* > *α* branch of this same decision.

One limit should be stated plainly. Which members compose the departing set, and how a parent group away from 2*N*\* resolves the competition among members to be in it rather than in the portion staying behind, is a coalition-formation problem that a per-capita treatment of interchangeable members does not solve. We use the sign of *α* − *φ* to fix which end of the *m* continuum departures concentrate at, which is what the distributional predictions below require, and we retain symmetric halving for the threshold results. The bargaining over composition is a natural target for future work, and it is where the heterogeneity and relatedness extensions taken up in the Discussion enter.

### 3.8 Fission costs and the realized fission threshold

The fission threshold derived above assumes that splitting is costless. In practice, fission carries direct costs: groups must physically separate, often losing access to part of their territory; social bonds are disrupted; coalitional alliances formed within the parent group may not survive the split; and newly separated daughter groups face heightened intergroup threat during the period of reduced size. Let *F* denote the per-capita cost of fission, borne by all members of both daughters at the moment of splitting. The realized fission threshold *N*_F_(*F*), defined in equation (5), is the smallest group size at which the post-fission payoff net of fission costs first exceeds the intact-group payoff:

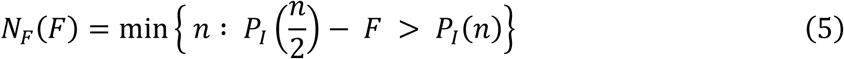

Because fission costs reduce the post-fission payoff, they delay the point at which splitting becomes worthwhile. The realized threshold *N*_F_(*F*) therefore lies at or above the costless threshold *N*_F_: equal when *F* = 0, and strictly above for any *F* > 0, increasing monotonically with *F*. At *F* = 0, the realized threshold equals the payoff-optimal threshold derived above. As *F* increases, the realized threshold rises toward the Nash equilibrium size — and as fission costs become prohibitively large, the realized threshold reaches the Nash equilibrium and the group never fissions. This limiting case recovers Sibly’s (1983) result: groups grow to the Nash equilibrium size under open-access conditions and stabilize there, without any fission regulatory mechanism operating.

Fission costs thus trace a continuum from the payoff-optimal threshold (costless fission) to the Nash equilibrium (prohibitively costly fission), with real systems occupying intermediate positions depending on the ecological and social costs of splitting. Notably, fission costs and slope asymmetry are independent parameters that can push in the same or opposite directions: a system with *β* > *α* (steep decline, low *N*_F_) and low fission costs may produce a lower realized threshold than a symmetric system with high fission costs, even though both have the same *N\**. The interaction of these two parameters determines the realized fission threshold and, through it, the time-averaged modal group size in a population.

One clarification is essential for what follows. Throughout the fission-only model, *F* is treated as a fixed parameter that determines the fission threshold *N*_F_(*F*). Groups fission when size crosses this threshold determined by *F*. This simplification isolates the effect of fission costs on the group size cycle. The fission-and-exclusion model, which follows, extends this framework by allowing fission to enter alongside admission and exclusion as a cost-bearing component of group regulation. This extension makes it possible to characterize when fission becomes the dominant response to group growth and when alternative mechanisms of regulation prevail.

### 3.9 Time-averaged modal group size under local joiner pools

With fission costs and slope asymmetry both in hand, we can now derive the time-averaged modal group size for a population of groups under local joiner pool conditions — the Case 1 regime in which each group draws from its own joiner stream independently. Under a constant joining rate of one new member per unit time, a group grows linearly from its post-fission starting size *N*_F_(*F*)/2 to its realized fission threshold *N*_F_(*F*). The time-averaged group size over this cycle is the midpoint of the growth range:

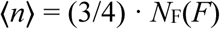

This can be expressed in terms of the underlying parameters. For the baseline case with symmetric slopes and no fission costs, ⟨*n*⟩ = (3/4)(4/3)*N\** = *N\**, the time average coincides exactly with the optimum, even though groups range from 2/3*N*\* to 4/3*N*\* at any given moment.

The direction of deviation from *N\** depends on the two parameters jointly. Fission costs always push ⟨*n*⟩ above *N\**: as *F* increases, *N*_F_(*F*) rises, and the entire growth cycle shifts upward. The group spends more time at supraoptimal sizes before the realized threshold triggers a split. Slope asymmetry pushes in a direction determined by the ratio *α*/*β*. When *α* > *β* (benefits accumulate faster than costs escalate), *N*_F_ is high and the group spends more time above *N\** than below it: ⟨*n*⟩ > *N\**. When *β* > *α* (costs escalate faster than benefits accumulate), *N*_F_ is low and the group spends more time below *N\** than above it: ⟨*n*⟩ < *N\**. These two parameters are additive in their directional effects on modal group size: when both push in the same direction — high fission costs together with *α* > *β* — modal size deviates strongly upward; when they oppose (e.g., steep decline but high fission costs), they partly counteract one another, with the dominant parameter determining the direction.

The full expression for time-averaged modal size under costless fission and asymmetric slopes is given in equation (6).

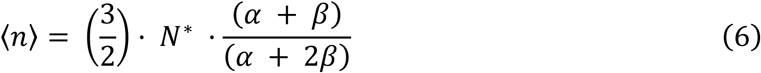

This exceeds *N\** when *α* > *β*, equals *N\** when *α* = *β*, and falls below *N\** when *β* > *α*. The slope ratio *α*/*β* is therefore the key structural parameter governing modal group size in local joiner pool systems under costless fission, with fission costs providing an independent upward shift that applies regardless of slope asymmetry.

### 3.10 Post-fission dynamics and the role of joiner pool structure

The payoff parameters *α*, *β*, and *F* established above determine the mechanics of the fission cycle for any single group: when it splits, how large its daughters are, and how long the group spends above versus below *N\** before splitting again. Two aspects of fission matter here and recur below. The first is the size at which a group fissions, which the threshold conditions above treat as symmetric halving, the tractable baseline retained throughout for *N*_F_, *N*_AF_, and *n*_EF_. The second is its geometry: how the parent’s mass divides. Symmetric halving is only the special case; real fissions extend to the asymmetric, in which a small group dissociates from a crowded group, while the bulk persists. This too follows from the individual rationality that governs joining, once one notes that *α* and *β* are merely group averages: a group is a set of individuals in shifting dyadic relationships, and the costs of crowding fall unevenly. As such, a member bearing a private cost well above the mean can rationally leave, alone if need be, even when the group-average payoff looks tolerable. We hold split geometry at symmetric halving for the threshold results below and let it vary when we turn to the population distribution, where it proves decisive. These parameters, however, say nothing about what happens to those daughters in a landscape that may contain other groups competing for the same prospective joiners. That question is governed by a second, independent dimension of the model: the structure of the joiner pool. These two dimensions (payoff parameters and joiner pool structure) interact to produce the full range of modal group size outcomes that the fission-only model predicts. The payoff parameters set the size and duration of the fission cycle; the joiner pool structure determines whether post-fission daughters each run their own cycle autonomously or compete for the same joiners, and therefore whether the population distribution of group sizes is driven primarily by within-group fission-regrowth dynamics or by differential daughter growth across a shared landscape. We consider three cases, each internally consistent with the payoff assumptions above and each producing a qualitatively distinct population-level outcome.

After fission at *N*_F_(*F*), two daughter groups exist, both starting at *N*_F_(*F*)/2 — below the nominal optimum and therefore attractive to prospective joiners under the entry condition.

#### 3.10.1 Case 1: Local joiner pools, independent growth

When prospective joiners arrive at each group independently from local catchments (e.g., because groups occupy separate territories, are rarely encountered simultaneously, or because joiners assess only one group at a time), post-fission daughters each draw from their own joiner streams. Both daughters grow autonomously and the population-level modal size is determined by the cycle-averaged formula derived above. Under symmetric slopes and costless fission, modal size is *N\**; under asymmetric slopes or positive fission costs, it deviates in the directions described. The population at any snapshot contains groups distributed across the growth cycle from *N*_F_(*F*)/2 to *N*_F_(*F*), with the time average as the central tendency. This is the appropriate model for relatively isolated groups or taxa in which prospective joiners encounter groups sequentially rather than comparatively.

#### 3.10.2 Case 2: Global joiner pool, asymmetric fission (the bimodal mechanism)

When prospective joiners can assess and compare multiple groups across a shared landscape, Ideal Free Distribution logic applies: joiners prefer the group offering the highest post-entry per-capita return. Under symmetric fission this does not strand the smaller daughter for long — joiners are pulled to whichever group is climbing toward the optimum, so both daughters pile up just below the fission threshold, leaving the standing distribution supra-optimal rather than sub-optimal. Permanent stranding of daughter groups results only at the costless symmetric limit (*F* = 0), where the unstranded, growing daughter reaches the fission threshold exactly as its post-entry payoff would fall to the smaller daughter — instructive, but biologically unexpected. The generic, robust route to sub-optimal sizes is asymmetric fission: more commonly, one or more members dissociate from a crowded group while the bulk persists. The per-capita payoff that defines crowding is a group average, but the underlying costs of conflict are not — a group is a set of individuals in shifting dyadic relationships, and a member bearing a private cost well above the mean does better by leaving, alone if need be, than by staying. A small daughter formed this way sits far below *N*\* and is the least attractive option in the pool for joiners; it lingers at small size while large groups keep recruiting and drift to or beyond *N*\*, producing a bimodal standing distribution: supra-optimal large groups coexisting with a sub-optimal mode of small daughters.

This requires that the departing member found a new group rather than join an existing one, and the requirement is not innocuous: the same global pool that lets joiners compare groups is available to a leaver, who would otherwise join the best group on offer rather than sit alone. The low mode therefore forms only where founding is the leaver’s realistic option, because no acceptable group exists for them to join. Where a leaver has available joining options instead, the same dissociation regulates size continuously and mints no small daughters, and the standing distribution is unimodal near the optimum. What decides whether the bimodality appears is therefore whether the large groups reach the Nash size. A joiner drawn from the global pool has the solitary return R1 as its alternative, so groups grow until a further entrant would gain no more than R1, which is the Nash size. At that point a small daughter of size *m* offers its own members more than joining a Nash-sized group would (the daughter returns the rising-limb payoff, the Nash group returns only R1), so the daughter persists rather than dissolving. Where the pool is instead closed, so that a prospective joiner is already a member of another group and compares that group rather than the solitary return, groups equilibrate near the optimum, no group approaches Nash, and any small daughter’s members do better rejoining: the low mode dissolves and the distribution is unimodal. The bimodal case is thus the open global pool.

Mechanically this is a coagulation–fragmentation balance (Lerch and Abbott, 2024; Ma et al., 2011): asymmetric fission continually produces small group fragments while joining allows fragments to coagulate back into larger groups, so any one small group is transient even though the low group size mode it populates is permanent and continually refilled. Departures concentrate at the smallest sizes here because this case is the *φ* > *α* branch of the departure decision derived above, in which each member added to a departing set costs more to bring along than the rising branch returns. Where *φ* < *α* the same decision produces collective departure and mints no small daughters at all, which is why the bimodal signature belongs to this branch specifically rather than to fission generally. The branch is load-bearing in a second way. Under strictly interchangeable members a lone departure becomes rational only above the Nash size (Appendix E), so the heterogeneity invoked just above, that the costs of conflict fall unevenly and some member bears a private cost well above the mean, is what makes the low mode form at all rather than a refinement of it. Given that heterogeneity, the height of the low mode is set by two observable quantities, the dissociation rate and the residence time of small groups, largely independent of slope asymmetry and fission cost, which instead govern the position of the supra-optimal mode as in Case 1 (Figure 2). The pattern reproduces the coexistence of large groups with lone individuals and small groups characteristic of fission-fusion societies across taxa (Couzin and Laidre 2009) and well documented among primates (Aureli, et al. 2008; Lehmann and Boesch 2004).

**Figure 1.**
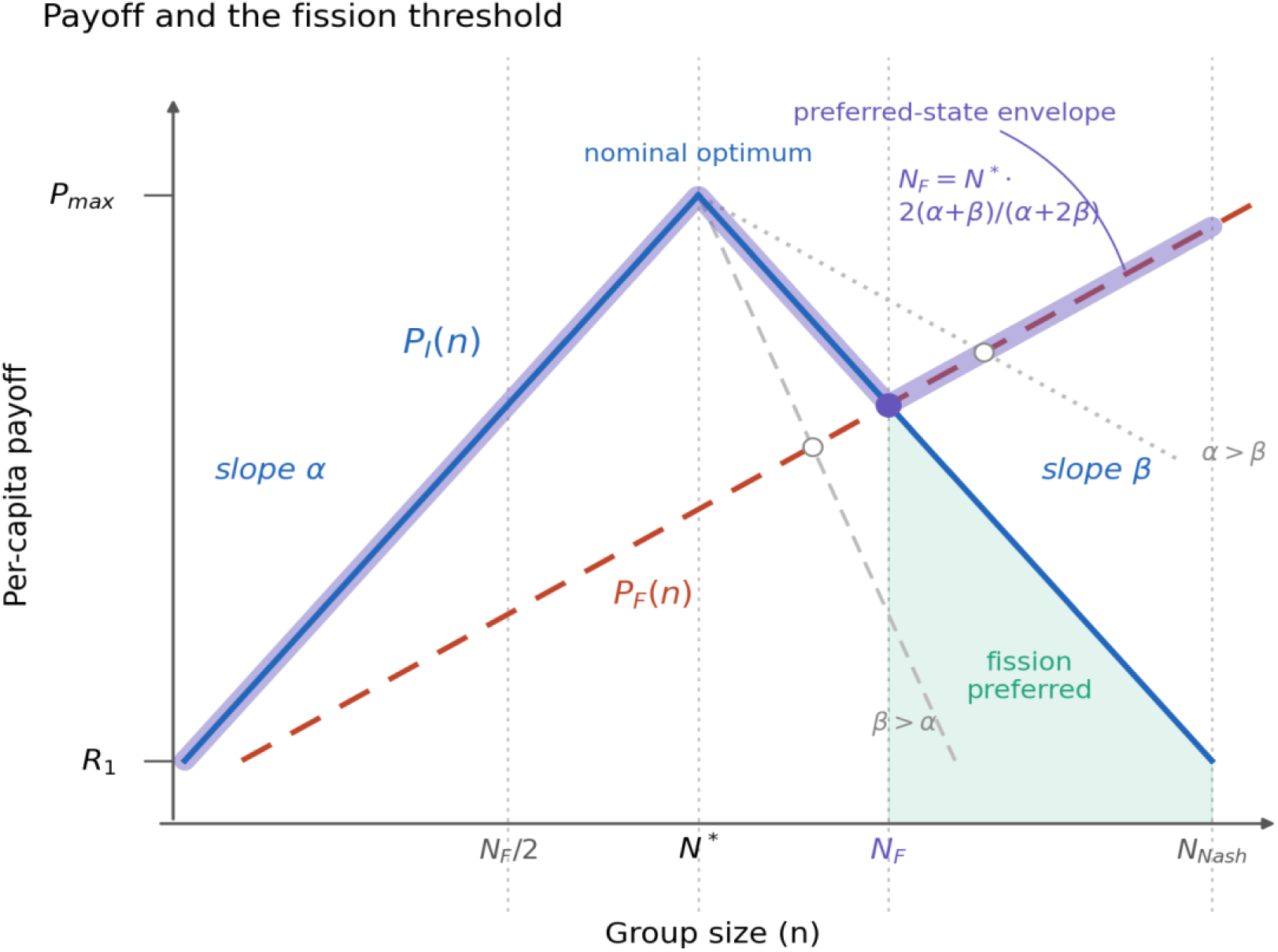
per-capita payoff as a function of group size under the fission-only model, shown for three cases: symmetric slopes (*α* = *β*, solid lines), steeper decline (*β* > *α*, dashed), and shallower decline (*α* > *β*, dotted). In each case the payoff function consists of two line segments meeting at the nominal optimum *N\**. The fission threshold *N_F_* (i.e., the group size at which splitting into two equal daughters first yields a higher per-capita return than remaining intact) shifts leftward when the decline slope is steep and rightward when it is shallow. Post-fission daughter size (= *N_F_*/2) and time-averaged group size over the fission-regrowth cycle shift accordingly. The preferred-state envelope (thick line) follows whichever payoff (intact or post-fission) is higher at each group size.

**Figure 2.**
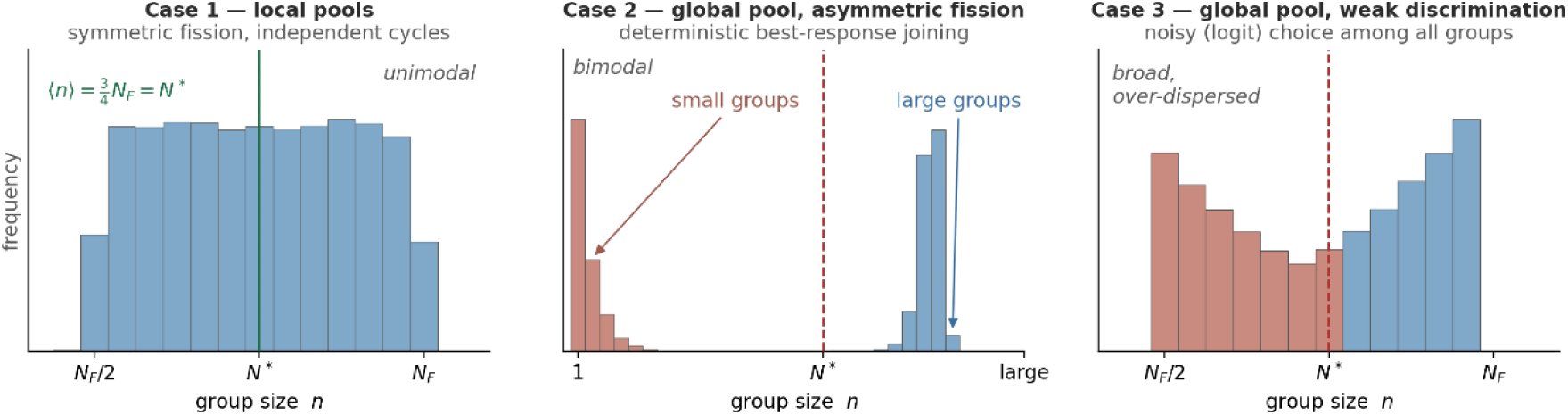
Predicted standing distributions of group size under the fission-only model, for three combinations of joiner-pool structure and fission geometry (optimum *N\** dashed; symmetric, costless baseline). In Case 1 (local pools, symmetric fission), independent fission–regrowth cycles give a unimodal distribution centered on the cycle-averaged size ⟨*n*⟩ = (3/4)*N_F_*, which equals *N\** in this baseline. In Case 2 (unequal access to the joiner pool), a small number of members leave larger, crowded groups while large groups continue to recruit under deterministic best-response joining, yielding a bimodal distribution: supra-optimal large groups together with a sub-optimal mode of small groups (orange). The small mode is continually refilled by asymmetric fission, and its height is set by the dissociation rate and the residence time of groups; a coordination cost that rises with the size of the departing group concentrates departures at the smallest sizes. In Case 3 (global pool, weak discrimination), stochastic (logit) choice among all groups yields a broad, over-dispersed distribution spanning sub- and supra-optimal sizes. Distributions are simulations of the corresponding dynamics; see <u>Zenodo repository</u> for the accompanying code (Figure2.py). These are the distributions the mechanism produces at a single optimum. An observed distribution pools this across places whose optima differ, and where the resource base varies appreciably from place to place that variation dominates the spread of observed sizes: the mechanism above shifts sizes by some tens of per cent around N*, while N* itself may differ several-fold between a settlement at a mission or a market and one in the interior. The distributions below are therefore predictions about deviation from a local optimum rather than about the shape of a pooled census, and the comparisons that isolate them hold the optimum fixed, either within a group over time or across groups matched on resource base.

#### 3.10.3 Case 3: Global joiner pool, stochastic choice under weak discrimination

Joiners may be unable to reliably distinguish small payoff differences between available groups. After fission, daughters may differ only slightly in size and per-capita return; those differences may be too subtle to detect from a distance, too slow to manifest in observable cues, or too easily swamped by proximity and chance encounter. Where this is so, joining is probabilistic rather than strictly payoff-maximizing. Both daughters receive joiners but not in equal proportion: the larger daughter is more likely to be preferred in any given encounter, so it grows faster in expectation. The strength of this amplification depends on the ratio of payoff difference to assessment noise. When daughters are nearly equal, noise dominates and joiners distribute close to randomly; as size differences grow, joiner discrimination becomes more decisive and the dynamics approach Case 2.

The Case 1–2 continuum can be formalized with a logit choice rule in which a noise parameter σ indexes joiner discrimination precision; the full treatment including equilibrium dynamics and empirical estimation of 1/σ is given in Appendix D.

### 3.11 Central result of the fission-only model

The fission-only model maps modal group size onto three independent sets of parameters: the payoff function parameters (*α*, *β*, and fission cost *F*) that govern the fission cycle, the departure coordination cost (*d* and *φ*) that governs the geometry of each split through the sign of *α* − *φ*, and the joiner pool structure (local versus global, and discrimination precision) that governs how post-fission daughters compete for joiners. The two sets interact to produce a parameter space with distinct regions corresponding to different modal group sizes.

The direction of modal group size deviation from *N\** is system-specific, determined by identifiable ecological and behavioral parameters. Groups in systems dominated by rapidly escalating internal costs (*β* > *α*) and global joiner pools should show strong suboptimal modal sizes; groups in systems with slowly accumulating costs (*α* > *β*), high fission costs, and local joiner pools should show supraoptimal modal sizes. The same payoff-function logic, the same individual rationality, and the same fission mechanism produce opposite directional predictions depending on parameters that are in principle observable. The result is a comparative empirical program: predict the direction in which any given population’s modal size deviates from its nominal optimum from a small set of ecological and behavioral parameters that can be measured with field data: α/β slope asymmetry, joiner pool structure, and fission cost.

A further implication concerns within-population variance in group size. Case 1 predicts a unimodal distribution of group sizes across the fission-regrowth cycle, centered at ⟨*n*⟩ with spread determined by *N*_F_(*F*) – *N*_F_(*F*)/2. Case 2 predicts a bimodal distribution: a sub-optimal mode of small groups alongside a supra-optimal mode of large groups, with the small mode’s height set by the dissociation rate and group residence time. Case 3 predicts a broad, over-dispersed distribution spanning sub- and supra-optimal sizes. These distributional signatures are empirically distinguishable and provide a test of which regime a population is in, independently of the mean modal size. Figure 2 shows the three predicted signatures side by side.

## 4. The fission-and-exclusion model and the viable exclusion condition

### 4.1 Setup

The fission-only model, introduced in Section 3, treats group maintenance as a passive cost environment: the declining slope *β* of the payoff function captures the ambient effects of within-group competition, conflict, crowding, and disease as costs that accumulate automatically with group size, regardless of what incumbents do. Within the fission-only model, fission cost *F* is also treated passively — as a threshold parameter that shifts the realized fission point *N*_F_(*F*). The fission-and-exclusion model promotes both the cost of resisting unwanted joiners and the decision to fission from passive to active. Incumbent groups can now respond to dilution by attempting coordinated exclusion at a real per-capita cost, and the decision to fission is itself weighed against exclusion at every step rather than triggered automatically at a threshold. The model now contains three genuine decision costs operating at every joiner arrival: the cost of admitting (a per-capita payoff loss *β* at sizes above *N\**), the cost of excluding (*C*_E_(*n*), a coordination cost rising with group size), and the cost of fissioning (*F*, the one-time per-capita penalty of splitting). Incumbents compare these three costs and act on the smallest.

### 4.2 The three-decision framework

When a prospective joiner arrives at a group of size *n*, the per-joiner decision is whether to admit or exclude that individual. Running underneath this binary choice is a group-level question that does not concern any single joiner: whether cumulative dilution has made splitting the whole group preferable to continuing under the better policy: admit or exclude. We write all three resulting payoffs on a common per-capita scale so that the two tiers can be compared directly, but they are not three interchangeable responses to one arrival — admit and exclude act on the prospective group joiner, whereas fission is a threshold the group monitors as it grows:

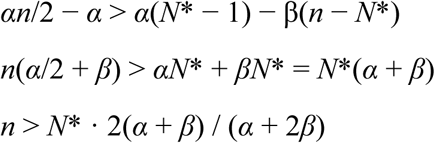

Admit allows the group to grow by one member; all incumbents and the newcomer share the diluted payoff *P*_I_(*n* + 1). Exclude maintains size at *n*, but incumbents collectively bear the per-capita cost *C*_E_(*n*) of organizing resistance. Fission splits the group equally into two daughters of size *n*/2 and imposes the per-capita fission cost *F* on every member. The optimal decision at any group size is whichever of *W*_A_, *W*_E_, and *W*_F_ is largest. As *n* varies, the maximum can shift among the three options, partitioning the size range into zones of preferred action. Precisely, the model is a two-tier hierarchy. At each arrival, incumbents first resolve the per-joiner contest between admission and exclusion, comparing *W*_A_ with *W*_E_; separately, they track whether growth has carried the group to the size at which fission first dominates the better of the two per-joiner options, i.e. whether *W*_F_ exceeds max(*W*_A_, *W*_E_). That second comparison is a threshold test of exactly the form used in Section 3’s fission-only model, with the admit-or-exclude winner replacing intact membership as the baseline.

The decision rule is one-step and local: at each joiner arrival, the group evaluates the three payoffs at its current size and acts on the maximum. The cumulative trajectory of a group across time emerges from repeated application of this rule as joiners arrive. We assume a constant arrival rate of one prospective joiner per unit time, as in the fission-only model.

### 4.3 Exclusion cost as a coordination problem

Exclusion is a public good: when it succeeds, every incumbent enjoys the smaller, less diluted group, whether or not they helped achieve it. Exclusion requires effort (e.g., monitoring, signaling, aggression, or coalition maintenance) that is often coordinated and borne privately by those who do the excluding. As group size grows, the number of incumbents whose can be coordinated around exclusion increases, and the temptation to free-ride on others’ effort grows correspondingly. The per-capita cost of exclusion rises with group size as a result.

We capture this with a simple linear cost function:

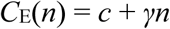

The *γn* term in *C*_E_(*n*) represents the expected cost of a coordination attempt that succeeds with a size-declining probability; the volunteer’s-dilemma micro-foundation is given in Appendix C. Together, *c* and *γ* specify how the second-order collective-action problem of exclusion translates into the cost paid by each willing excluder in a group of size *n*. Low *c* and low *γ* describe a system where coordination is cheap and free-riding is mild — small, well-organized groups or systems with clear sanctioning mechanisms. High *c* or high *γ* describe systems where coordination is costly or free-riding dominates as size grows. That larger groups secure collective goods less reliably, as individual willingness to contribute falls with size, is well-established (Archetti, 2009). The link between failed within-group coordination and fission is not a hypothetical construct: free-riding in coordinated exclusion is an experimentally documented driver of fission in human task groups (Hart and Van Vugt, 2006), where groups subject to within-group free-riding split along their pre-existing internal divisions.

The viable exclusion condition in equation (2) follows from comparing *W*_A_(*N\**) with *W*_E_(*N\**); the derivation is given in Appendix B. Figure 3 plots *W_A_*, *W_E_*, and *W_F_* against group size for both regimes, showing the group locking at *N*\* when exclusion is viable and fissioning at *N*_AF_ when it is not.

**Figure 3.**
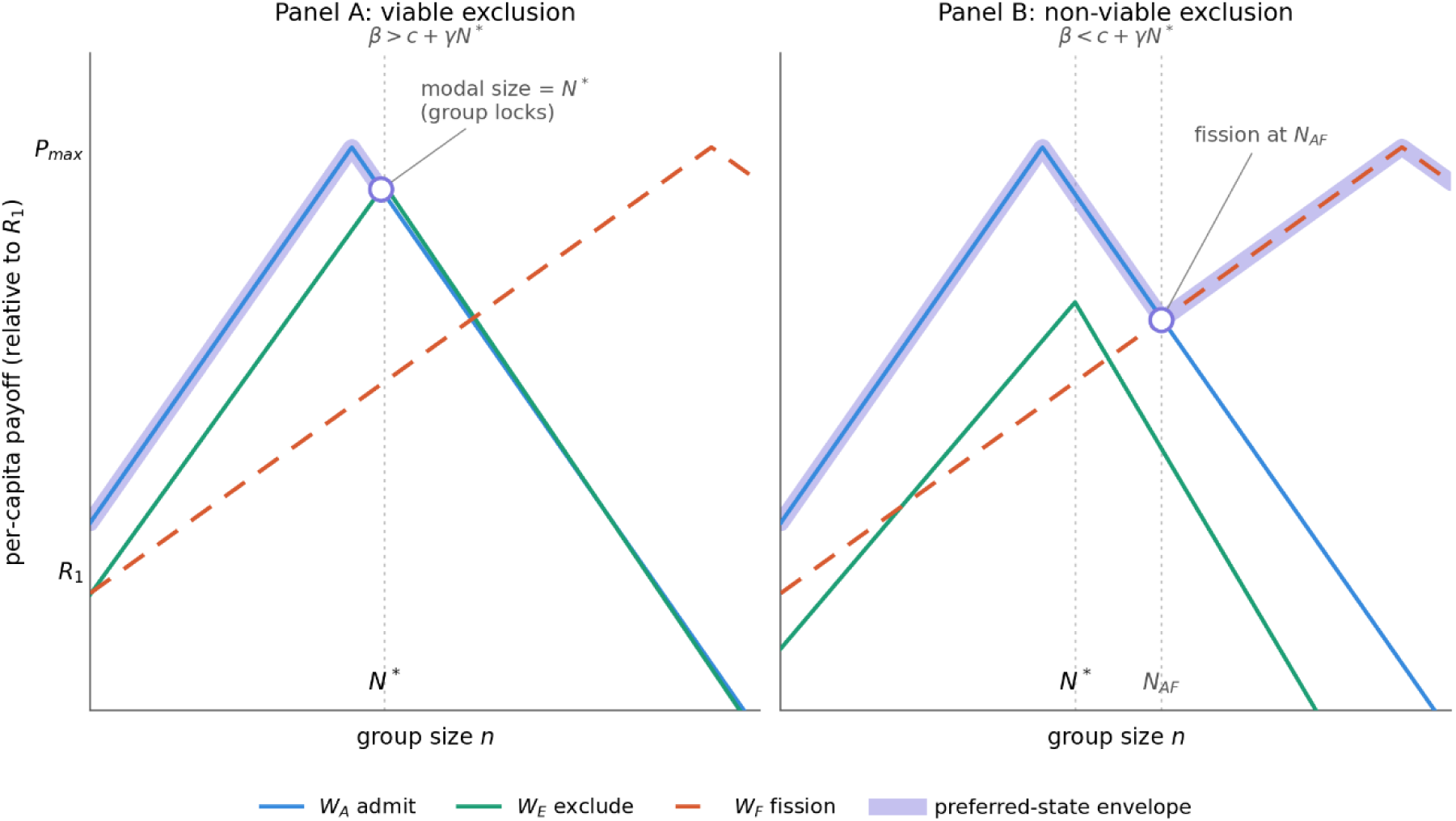
Drawn throughout under the symmetric-halving convention retained for the threshold results, so that *W_F_* is a single per-capita quantity and each daughter is *n*/2. Panel A: the exclusion viability condition depends only on comparing *W_E_* and *W_A_* at *N*\*; fission geometry is irrelevant. Panel B: *N_AF_* is the collective-departure branch; if the departure coordination slope exceeds the rising payoff slope, the group sheds members one at a time and drifts to the Nash size without a discrete split. The three decision payoffs *W_A_*(*n*) (admit), *W_E_*(*n*) (exclude), and *W_F_*(*n*) (fission) are plotted as functions of group size in two parameter regimes. The preferred-state envelope (the payoff-maximizing option at each *n*) is traced by whichever of *W_A_*, *W_E_*, and *W_F_* is highest. Panel A: *W_E_* crosses above *W_A_* at *N*\* and the preferred-state envelope switches from admit to exclude. Under sequential growth, the group reaches *N*\* and locks there indefinitely; no further joiners are admitted and fission is suppressed. Panel B: *W_E_* lies below *W_A_* throughout the relevant range and never enters the envelope. The group admits past *N*\* until the admit-fission crossing at *N_AF_*, where *W_F_* rises above *W_A_* and the envelope switches to fission; the fission-only dynamics are recovered. The transition between the two regimes is sharp at the boundary *β* = *c* + *γN*\*.

### 4.4 Two regimes of modal group size

The viability condition partitions parameter space into two regimes, each yielding a qualitatively distinct prediction about modal group size.

#### 4.4.1 Non-viable exclusion (β < c + γN*)

When exclusion is too expensive to beat admission at *N\**, the group never excludes. Joiners are admitted past *N\** until fission becomes preferred over admission. The admit-fission crossing *N*_AF_ is the smallest *n* at which *W*_F_ > *W*_A_. Solving:

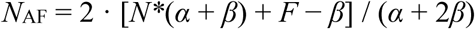

When *F* = 0, this is the costless fission threshold *N*_F_ from the fission-only model, up to a small discrete-step correction (because admission moves the group by one whereas fission halves it). For positive *F*, *N*_AF_ shifts upward as in the fission-only model. Within this regime, the introduction of exclusion as an option changes nothing: *W*_E_ is never on the envelope, the policy is identical to the fission-only model’s admit-only-then-fission policy, and modal group size is the value averaged over the fission-regrowth cycle (weighted by how many joiner-arrivals the group experiences at each size). The fission-only model’s full case structure (i.e., local pools, global pool with preference, weak discrimination) applies unchanged.

#### 4.4.2 Viable exclusion (β > c + γN*)

When exclusion is sufficiently cheap, *W*_E_ rises above *W*_A_ at *N\**. A group growing from below first encounters this crossing at *n* = *N\**: the joiner arriving when the group has just reached the nominal optimum is excluded rather than admitted. The group does not grow further. Under steady joiner arrival, the group remains at *N\** indefinitely, excluding every joiner that arrives. Modal group size in the viable regime is:

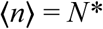

Fission never occurs in steady state. The group locks at *N\** before it has any opportunity to reach the fission threshold, and the fission decision is therefore never invoked under sequential growth dynamics. For completeness, the size at which exclusion would give way to fission if the group were pushed there by some external perturbation (e.g., demographic shock, environmental disturbance, or predation) is given by the exclude-fission crossing:

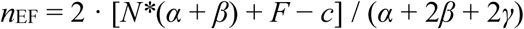

Under sequential growth from below *N\**, the group locks at *N\** and never approaches *n*_EF_. The exclude-fission crossing therefore characterizes a hypothetical policy boundary rather than a realized group size, and matters only when groups are placed above *N\** by exogenous factors, for example, historical fission events that occurred before exclusion became viable, environmental shocks, or internal birth rates that outpace the rate at which exclusion can repel them.

#### 4.4.3 The sharp regime transition

The shift between regimes is discontinuous: infinitesimal changes in *c*, *γ*, or *β* across the boundary *β* = *c* + *γN\** qualitatively reorganize the population dynamics. This has no counterpart in Section (3)’s fission-only model, where modal size varies continuously with payoff parameters; here, viable exclusion creates a sharp boundary across which dynamics restructure abruptly.

### 4.5 Joiner pool structure within the two regimes

The joiner pool structure (i.e., local pools, global pool with strong preference, and global pool with weak discrimination) interacts with the regime to determine population-level group size distributions. Within each case, the population dynamics depend on whether exclusion is viable, and the resulting distributions differ qualitatively across the six combinations, which Table 1 summarizes.

**Table 1.** Steady-state group-size distributions in the two exclusion regimes for each joiner-pool case. *N\** is the nominal optimum, *N*_AF_ the admit–fission crossing, and *n*_EF_ the exclude–fission crossing. Non-viable exclusion (*β* < *c* + *γN\**) reproduces the fission-only dynamics; viable exclusion (*β* > *c* + *γN\**) pins any group that can grow by accretion at *N\** and suppresses fission in steady state.

| Joiner-pool case | Non-viable exclusion<br>($\beta < c + \gamma N^*$) | Viable exclusion<br>($\beta > c + \gamma N^*$) |
| --- | --- | --- |
| **Case 1. Local pools (independent growth)** | Fission-only dynamics: each group cycles between $N_{AF}/2$ and $N_{AF}$. Unimodal; time-averaged modal size $(3/4)N_{AF}$, shifted by slope asymmetry and fission cost. Fission recurrent. | Every group grows to $N^*$ and locks. Unimodal at $N^*$ exactly. Fission absent in steady state (at most one transient split for groups initialized above $n_{EF}$). |
| **Case 2. Global pool, asymmetric fission (bimodal)** | Fission in crowded groups produces small daughter groups while large daughter groups cycle toward $N_{AF}$; small daughter groups sit below $N^*$. Bimodal: a sub-optimal mode of small groups below $N^*$ plus a cycling mode near $N_{AF}$. Small daughters continually minted; fission recurrent. | Larger daughter accretes to $N^*$ and locks; pre-existing daughter groups remain below $N^*$. Stable bimodal: one mode pinned at $N^*$, one small-daughter mode below $N^*$. No further fission; small-daughter fraction set by pre-viability history. |
| **Case 3. Global pool, weak discrimination (stochastic choice)** | Both daughters recruit, the larger faster. Modal size near $N^*$ with bimodal transients, increasingly left-skewed as discrimination sharpens. Fission recurrent. | Both daughters accrete and lock at $N^*$. Unimodal at $N^*$ at equilibrium (transient bimodality); slower approach than Case 1. No further fission. |

Two regularities cut across Table 1. First, viable exclusion collapses the differences among joiner-pool cases: any group that can grow by accretion is pinned at *N\**, so the three cases converge on a mode at *N\** (Case 2 retaining only a residual small-daughter mode inherited from fission events that predate viability). Second, the viability boundary toggles fission frequency: fission is recurrent throughout the non-viable column and absent from the steady state of the viable column. Joiner-pool structure therefore shapes the distribution only when exclusion is non-viable; once exclusion becomes viable it overrides that structure for every group not stranded below *N\**.

### 4.6 Central result of the fission-and-exclusion model

The fission-and-exclusion model maps modal group size onto seven independent parameters: the payoff function parameters *α* and *β* (from the fission-only model), the fission cost *F* and the departure coordination parameters *d* and *φ* (from the fission-only model), the exclusion cost parameters *c* and *γ* (from the fission-and-exclusion model), and the joiner pool structure (also from the fission-only model). The five-parameter space is partitioned by a single inequality, the viable exclusion condition in equation (2), into two regimes with qualitatively distinct dynamics.

Three empirically tractable predictions follow. First, the direction of modal group size deviation from *N\** is determined by the viability condition: systems satisfying equation (2) should show modal sizes at *N\** regardless of the joiner pool case, while systems violating it should show the case-specific deviations. Second, fission frequency drops to zero across the viability boundary: systems just inside the viable regime should show effectively no fission events in steady state, while systems just outside should show recurrent fission cycles. Third, the bimodal distribution of Case 2 persists in both regimes but its larger mode shifts from a cycling time-average (non-viable) to a pinned value at *N\** (viable), and the minting of new sub-optimal daughters halts when the system enters the viable regime.

The viability condition itself is empirically tractable. The slope *β* of the declining payoff function can be estimated from the rate at which per-capita measures of fitness, foraging success, or reproductive output fall as observed group size rises above the nominal optimum (Borries et al., 2008). The exclusion cost parameters *c* and *γ* can be estimated from behavioral measures of resistance effort and from the rate at which that effort fails as group size increases. Comparative tests across taxa or social systems can then map populations onto either side of the viability boundary and compare predicted to observed group size distributions. The framework provides a quantitative criterion for predicting exclusion viability across systems.

## 5. Discussion

The framework’s empirical reach spans the major taxonomic groups in which fission-fusion dynamics have been studied, including the skewed and often bimodal group-size distributions documented across many taxa (Couzin and Laidre, 2009). The fission-only model and the fission-and-exclusion model together unify predictions that have been derived independently across fishes (Buston, 2003; Pitcher, 1986), social insects (Grinsted and Field, 2018a), cooperatively breeding birds (Sturrock et al., 2022), rodents (Waterman, 2002), primates (Borries et al., 2008; Markham et al., 2015; Sandel et al., 2026a; Wakeford and Cords, 2025), and human forager groups (Walker and Hill, 2014). We focus the comparative discussion that follows on chimpanzee (*Pan troglodytes*) and human forager cases, where the empirical literature on group-size dynamics is most developed and where we can directly examine the framework’s predicted regime signatures. What unifies these cases methodologically is a single question, made tractable by the framework: how is the distribution of group sizes positioned and shaped relative to the nominal optimum, and why? Four measurable parameters do the work: the slope asymmetry *α*/*β* of the per-capita payoff function, the structure of the joiner pool, the per-capita cost of coordinated exclusion, and the fission cost. Each governs how readily one of the three modes of conflict resolution (admission, exclusion, and fission) becomes the dominant response in a given system. These cases also set up a prediction the framework makes that no prior account does: that the paired daughters of a single fission should diverge systematically in how readily they admit newcomers, a comparative prediction we develop below.

Among wild primates, the puzzle of modal group sizes that deviate systematically from per-capita optima has remained an active concern of the field (Markham and Gesquiere, 2017). Chimpanzees are a natural test case, because the size and composition of their parties deviate from per-capita optima and reorganize far more than food supply alone would predict — exactly the directional deviation the framework addresses. The framework treats two things as one process at different timescales: the everyday flux in which parties form, grow, shed members, and dissolve, and the rare permanent fission in which a whole community divides. The first is where the model’s machinery operates directly; the second is its limiting case, reached when the same conflict, accumulated over years, hardens party boundaries into a community split. Food sets the stage for party size: its distribution governs where females range and how contestable access to them becomes (Emery Thompson et al., 2007; Murray et al., 2007; Pusey et al., 1997), which is why party size responds more to how food is distributed than to its overall abundance, and why food’s influence wanes where food is plentiful (Lehmann and Boesch, 2004; Newton-Fisher et al., 2000). Protective benefits set the stage for community size: a larger community holds a larger territory and faces less pressure from neighbors, so between-group competition pushes community size upward and holds communities together above their nominal optimum (Lemoine et al., 2020; Wilson et al., 2014). But what regulates party size on that stage is the conflict over mating access that the resulting distribution of estrous females generates: their presence draws larger parties even as the competition it provokes among males can drive some apart, so party size reflects how that conflict is mediated, moment to moment, through who is admitted, tolerated, and driven out (Goodall 1986).

What our framework identifies is a balance between the conflict that accumulates within a group and the ease with which individuals resolve it by regrouping. The decisions the model formalizes are played out moment to moment at the party level, where individuals aggregate with a subgroup, are tolerated or not, and dissociate to join or form others. The base model’s fission into comparable daughters is one end of a continuum; at its other extreme, a single individual dissociates, sometimes drawing followers, to form a smaller party that may later re-fuse, an asymmetric and impermanent fission (Aureli, et al. 2008). Such dissociation is the joiner-pool dynamic running in real time: the flexible, arousal-sensitive party composition that field studies describe, featuring a male core whose membership shifts within and across days (Goodall, 1986; Kawanaka, 1984). The cost that regulates it is borne by the dissociating individual who forfeits access to contested resources and is exposed to threats that a larger party buffers (Boesch, 1991; Lehmann and Boesch, 2004). A daughter party formed this way can therefore inherit a lower per-capita payoff than its parent. Community size sets how readily the party level reabsorbs an individual who dissociates: small communities sustain larger, longer-lasting parties with closer association between the sexes, while large communities sustain smaller, shorter-lasting parties with weaker association, and beyond some point these parties stop exchanging members (Lehmann and Boesch, 2004). Permanent community fission is the rare limiting case of this same process, when mating-access conflict intensifies faster than the party level can absorb it, which is why such splits are separated by roughly 500 years (Langergraber et al., 2014). Ngogo, the largest community ever recorded at roughly 190 individuals, is the best-documented: association between two clusters progressively broke down until it split permanently into antagonistic factions (Sandel et al. 2026). Fission therefore forfeits this protective advantage: a daughter community holds a smaller territory and can, at the extreme, be destroyed by its neighbors, as the Kahama community was after splitting from Kasekela (Goodall, 1986; Wilson et al., 2014). The framework thus supplies a single account in which the admission, exclusion, and fission (dissociation) decisions that shape a party from hour to hour are the same decisions that, accumulated and hardened, determine whether a community holds together or divides, wherever such decisions are made.

Smith’s (1985) analysis of Inuit foragers was foundational in directing attention to conflict-mediated group size regulation. Since then, work across diverse forager populations has mapped how resource distribution, kinship structure, and sources of conflict shape group size and when groups fission, fuse, and reassemble (Hamilton et al., 2007; Hill et al., 2011). Among these, kinship and protection determine where group size sits before conflict regulates the variation around it. Kinship works through relatedness: shared production, alliance, and cooperation accrue along kin lines, so that denser kin networks and shared production, effectively raising α relative to β, settle at larger optima (Walker, 2014); ecological models predict these optima closely, as when Janssen and Hill (2014) derive an optimal 7–8 hunters for the Ache. Protection works through numbers: intergroup lethal violence was a regular feature of mobile forager life and tracked resource scarcity (Allen et al., 2016; Glowacki, 2025), so where neighboring groups are hostile, the advantage of greater numbers holds communities together above their within-group optimum, the human parallel to the between-group force in chimpanzees. On these foundations the same conflict-mediated regulation operates. Kinship sets the lines along which a community can divide, and what drives it to divide is internal conflict over leadership, alliance, and the distribution of shared production, in which mating and marriage are central rather than incidental. The durable pair bond is central to this, since who marries whom largely determines alliance membership, such that mating competition is embedded in broader social struggles over coalitional support and resources. When such conflict outgrows what kinship and marriage alliances can contain, the community fissions along kin lines. Walker and Hill’s (2014) cross-cultural review supports this. Their sample is drawn from semi-sedentary and sedentary village societies, mobile foragers being excluded because their groups reassemble continuously rather than splitting at discrete moments; across it, internal political conflict is the primary reported driver in 29 of 37 documented fissions and resource scarcity in only 8, and the two are often conflated, matching the prediction that fission resolves accumulated intragroup conflict. The best-documented cases tell one story in different registers. Disputes over women and leadership drove fissions among both the Yanomamö and the Xavante (Chagnon, 1979; Maybury-Lewis, 1967), jealousy and in-fighting split Kayapó villages seven times over (Verswijver, 1992), and at Old Orayvi in 1906, the most thoroughly studied fission on record, a Hopi village of over a thousand divided along factional lines that followed lineage membership (Whiteley, 2008). In each, what the village could no longer contain was conflict among its own members.

The fission-only and fission-and-exclusion models combine to produce a prediction about post-fission behavior that neither part anticipates alone. After fission, paired daughters face asymmetric incentives regarding admission of newcomers. The larger daughter — closer to *N*\* and preferred by joiners — has reason to become selectively restrictive: unchecked growth will move it to the fission threshold, so the benefits of coordinating exclusion are real and immediate. The smaller daughter faces the opposite incentive structure: it is below *N*\*, unattractive relative to its paired daughter, and liable to linger as a small, sub-optimal group. Admitting joiners freely — lowering effective entry costs, tolerating individuals it might otherwise exclude — is its best available strategy for growing back toward the optimum. The two daughters should therefore diverge in admission behavior in the period immediately following fission, with the larger becoming more restrictive and the smaller more permissive, even when they were drawn from the same parent group under identical ecological conditions. The prediction is empirically tractable: recently fissioned daughter groups of similar ecological quality should differ in their tolerance of newcomers, with the smaller daughter showing systematically lower effective entry barriers following the split. To our knowledge, no existing theory produces this specific prediction of divergent post-fission admission tolerance, and no empirical analysis tests for it; the framework therefore generates a tractable comparative question in fission-fusion research.

One assumption deserves to be stated as a scope condition rather than left implicit, because it is measurable and it marks where this account stops. The per-capita payoff P_I(n) is the average return to membership, and it is a true description of what any particular member receives only where returns are levelled. Where they are not, the average describes nobody, and it describes least well the members who are subsidizing the rest. So the model’s reach is the condition that realized dispersion of returns around their mean is small, with dispersion of holdings within a group serving as an observable estimate of it. This matters beyond bookkeeping. Departure, in this account, is a comparison between what membership returns and what leaving is worth, and both quantities become member-specific as returns disperse: the members who would leave first are those whose realized return has fallen furthest below the average and whose alternatives are best. That is a model of composition rather than of size, and it is a different paper. The present results describe size, and they hold where levelling holds. Several further extensions are tractable but lie beyond the scope of this paper. The framework treats joiner arrival as a constant-rate process with one prospective joiner at a time. Where many joiners arrive simultaneously and compete for entry, the bargaining structure inverts: incumbent groups gain power over candidates, the cost of selectivity shifts onto applicants, and the appropriate model becomes a matching market with two-sided preferences (Gale and Shapley, 1962; Noë and Hammerstein, 1994). The framework also treats the joiner pool as homogeneous; in practice joiners vary along multiple dimensions — spatial proximity, kinship, age and sex composition, skill and knowledge, behavioral profile (Bowler and Benton, 2005; Schniter et al., 2018, 2015; Sih et al., 2004; Wakeford and Cords, 2025; Widdig et al., 2006)— and daughter groups facing heterogeneous pools should differentiate to occupy distinct niches, a Hotelling-style dynamic that adds a second axis of post-fission divergence orthogonal to the size-based axis above. The relatedness extension introduced by Smith (1985) folds naturally into the framework as a reweighting of per-capita payoffs by within-group relatedness, though we have set that step aside here to isolate the structural results. The framework also defines a comparative test across the viable exclusion boundary that we do not run here: field estimation of the exclusion-cost parameters *c* and *γ* would place real systems on one side or the other, a tractable next step the model makes precise. Finally, the framework does not address how groups acquire the capacity for coordinated exclusion in the first place, nor does it treat fission as the extended social process documented in long-term studies of permanent splits (Feder et al. 2025; Sandel et al. 2026); these are also natural targets for future work.

## 6. Conclusion

Observed group sizes emerge from conflict-mediated regulation through admission, exclusion, and fission decisions. Whether a group sits below, at, or above its nominal optimum depends on a small set of measurable parameters: the slope asymmetry of the per-capita payoff function, the structure of the joiner pool, and the cost of coordinated exclusion. A single inequality, the viable exclusion condition, separates two qualitatively distinct dynamic regimes. The framework provides a mechanistic account of why, within populations, observed group sizes diverge systematically from the sizes that would maximize individual payoffs, and it applies across taxa.

## Funding

This research received no specific grant from any funding agency in the public, commercial, or not-for-profit sectors.

## Conflict of Interest

The author declares no conflict of interest.

## Data and Code Accessibility

This paper presents theoretical models only; no empirical data were collected or analyzed. All code used to produce the figures and simulations in this paper is archived in a public repository at Zenodo: https://doi.org/10.5281/zenodo.20358982 The repository includes Figure1.py, Figure2.py, Figure3.py for generation of the manuscript figures, and a README.md describing all files, parameters, and usage instructions. The repository is publicly available (Schniter, 2026); the DOI is permanent and will not change.

## Appendix A: Derivation of the Fission Threshold

The fission threshold is defined as the smallest intact-group size at which splitting into two equal daughters yields a higher per-capita payoff than remaining intact. Starting from the piecewise linear payoff function:

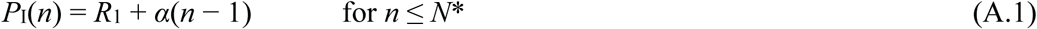

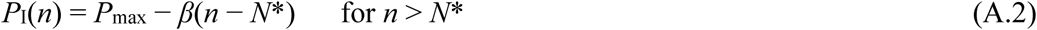

and noting that post-fission daughters begin below *N*\* so *P*_I_(*n*/2) = *R*_1_ + *α*(*n*/2 − 1), the threshold condition *P*_I_(*n*/2) > *P*_I_(*n*) on the declining branch becomes:

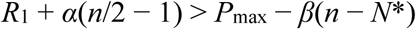

Substituting *P*_max_ = *R*_1_ + α(*N*\* − 1) and collecting terms in *n*:

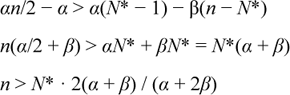

The fission threshold *N*_F_ is the smallest integer satisfying this inequality:

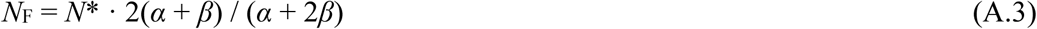

In the symmetric case (*α* = *β*) this simplifies to *N*_F_ = ⁴⁄₃*N*\*. Post-fission daughter size is *N*_F_/2 = *N*\*(*α* + *β*)/(*α* + 2*β*), equal to ⅔*N*\* when *α* = *β*. When fission carries a per-capita cost *F*, the realized threshold satisfies *P*_I_(*n*/2) − *F* > *P*_I_(*n*), shifting the threshold upward by 2*F*/(*α* + 2*β*), so it increases monotonically with *F* and approaches the Nash equilibrium size as *F* → ∞.

## Appendix B: Derivation of the Viable Exclusion Condition

Exclusion is viable if it dominates admission at *N*\*, the smallest size at which adding a member reduces per-capita payoff. The relevant payoffs evaluated at *N*\* are:

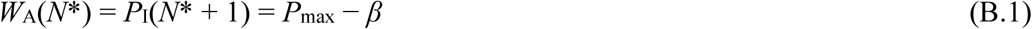

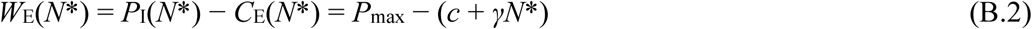

Exclude beats admit at *N*\* if and only if *W*_E_(*N*\*) > *W*_A_(*N*\*), i.e.:

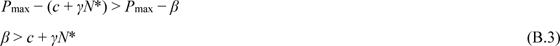

This is the viable exclusion condition. For *n* < *N*\*, admission always raises per-capita payoff so *W*_A_(*n*) > *W*_E_(*n*) always holds; the condition binds only at *N*\* and above. For *n* > *N*\*, *C*_E_(*n*) = *c* + *γn* continues to rise relative to the fixed cost *β* per additional member, so if exclusion beats admission at *N*\*, it does so for all *n* ∈ [*N*\*, *N*_F_]. When *β* ≤ *c* + *γN*\*, exclusion never beats admission anywhere in the relevant range, and the group grows past *N*\* until fission is triggered.

The exclude–fission crossing *n*_EF,_ from Section 4.4.2, follows from equating the exclude and fission payoffs at a size above *N*\*. Writing *W*_E_(*n*) = *P*_I_(*n*) − (*c* + *γn*) and *W*_F_(*n*) = *P*_I_(*n*/2) − *F*, with the declining-branch and rising-branch forms of *P*_I_ used in Appendix A, the crossing *W*_E_ = *W*_F_ gives, after substituting *P*_max_ = *R*_1_ + *α*(*N*\* − 1) and collecting terms in *n*:

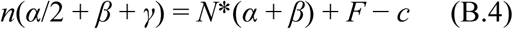

so that

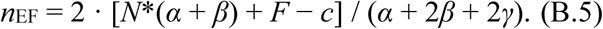

The *γ* term, absent from the admit–fission crossing *N*_AF_, enters because the per-capita exclusion cost rises with size whereas the per-entry admission cost *β* does not; it is what places *n*_EF_ below the size the same group would reach under admission alone. As emphasized in Section 4.4.2, a viable-exclusion group growing sequentially from below *N*\* locks at *N*\* and never approaches *n*_EF_, which therefore characterizes a policy boundary relevant only to groups displaced above *N*\* by exogenous factors.

## Appendix C: Volunteer’s-Dilemma Micro-Foundation for *C*_E_(*n*)

The exclusion cost function *C*_E_(*n*) = *c* + *γn* is parameterized so that *c* captures the baseline per-capita effort of attempting exclusion and *γn* captures the size-dependent rise in expected cost. The *γn* term is not a claim that the physical effort of a successful exclusion attempt grows with group size; conditional on success, per-capita effort is roughly constant. What changes with *n* is the probability that coordination succeeds.

Following the volunteer’s-dilemma framework (Archetti 2009), the chance that the required contributors actually volunteer declines faster than the number of potential contributors rises as group size grows. The expected cost to an incumbent — per-attempt effort divided by the probability of successful coordination — therefore rises with *n*. Writing this expected cost as *e* / *p*(*n*), where *e* is the per-attempt effort and *p*(*n*) is the size-dependent coordination success probability, the *γn* term approximates the contribution of declining *p*(*n*) to the expected cost over the relevant range. The linear approximation is tractable and preserves the qualitative monotonicity that drives the viable exclusion boundary; the results hold for any monotonically increasing *C*_E_(*n*) satisfying *C*_E_(1) = *c* > 0.

## Appendix D: Formal Model of Case 3 (Weak Joiner Discrimination)

Case 3 covers populations in which prospective joiners can perceive payoff differences between available daughters but do so imperfectly. We formalize this with a logit choice rule indexed by an assessment-noise parameter σ.

### The choice rule

Let the two post-fission daughters have current sizes *m_L_* (larger) and *m_S_* (smaller), with per-capita post-entry payoffs *P_I_*(*m_L_* + 1) and *P_I_*(*m_S_* + 1). A prospective joiner chooses the larger daughter with probability:

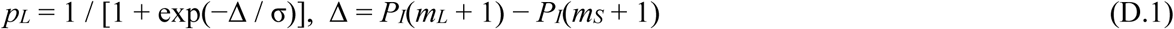

where Δ ≥ 0 is the post-entry payoff advantage of the larger daughter and σ > 0 is the assessment-noise scale. As σ → 0, *p_L_* → 1 whenever Δ > 0 and the rule collapses to deterministic preference for the larger daughter. As σ → ∞, *p_L_* → ½ and joiners split evenly, running independent Case-1-like cycles. Intermediate σ produces stochastic asymmetry: both daughters receive joiners, the larger with probability above ½.

### Gap dynamics

The expected change in the size gap per arrival is:

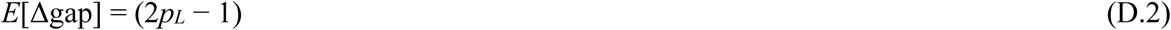

This is positive whenever Δ > 0 and increases with Δ/σ, so the gap widens in expectation at a rate that measures how strongly the population is pushed toward the asymmetric outcome. Slope asymmetry enters through Δ: daughters on the rising limb just after fission produce Δ ≈ α per unit size difference, so steep-decline systems (low *N_F_*, daughters far below *N*\*) push Case 3 toward Case 2, while shallow-decline systems (high *N_F_*, daughters near *N*\*) keep allocation near even.

### Empirical estimation

The discrimination strength 1/σ is estimable from observed joining decisions. Given binary choices between groups with independently estimated payoff differences Δ, σ is recovered by fitting the logit above — a standard discrete-choice estimation. The slope of the empirical choice probability against Δ near Δ = 0 identifies 1/σ. A population’s position on the Case 1–2 continuum is then a measured quantity, and the fit yields a test of whether joiners discriminate at all (σ finite) against the null of random affiliation (σ → ∞).

## Appendix E: Derivation of the Second Viability Condition

A departing set of size *m* leaving a group of size *n* > *N*\* bears the fission cost *F* and the departure coordination cost D(*m*) = *d* + *φm*. Because departing sets form at or below the optimum, the post-departure payoff to a member of the set is taken from the rising branch of *P*_I_:

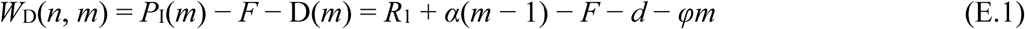

The parent size *n* does not appear, because a departing member’s payoff depends on the group the set forms rather than on the one it leaves. Differentiating with respect to *m*:

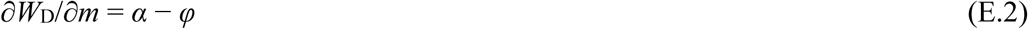

so *W*_D_ is monotone in *m* with the sign of *α* − *φ*, and the optimal size of the departing set lies at a corner of the admissible range 1 ≤ *m* ≤ *N*\*:

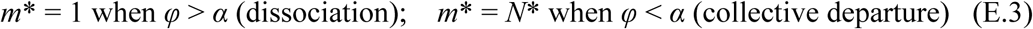

The upper limit is *N*\* rather than *n* − 1 because for *m* > *N*\* the payoff follows the declining branch and *W*_D_ falls at rate *β* + *φ*, so no departing set has reason to exceed the optimum. At *φ* = *α* the payoff is flat in *m* and the size of the departing set is not determined by these parameters.

The collective-departure branch. Departure occurs at the smallest *n* for which *W*_D_(*n*, *m*\*) exceeds the intact payoff *P*_I_(*n*) = *P*_max_ − *β*(*n* − *N*\*). With *m*\* = *N*\*, *W*_D_ = *P*_max_ − *F* − *d* − *φN*\*, and the condition gives

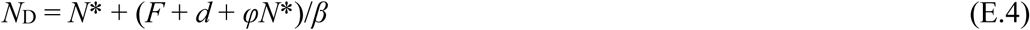

with a departing daughter of *N*\* and a remainder of *N*_D_ − *N*\* = (*F* + *d* + *φN*\*)/*β*. The split is even, and the symmetric halving of Appendix A is recovered, exactly when *F* + *d* + *φN*\* = *βN*\*, that is when the accumulated cost of leaving equals the payoff a member forgoes by staying at 2*N*\*. Away from that equality the same branch predicts an uneven split with one daughter at the optimum, a sharper prediction than halving and the one that daughter-size data can test.

The dissociation branch. With *m*\* = 1, *W*_D_(*n*, 1) = *R*_1_ − *F* − *d* − *φ*, which lies below *R*_1_ and therefore below the payoff of any intact group its members would rather belong to than not. Departure becomes rational only once *P*_I_(*n*) has fallen to that level, at

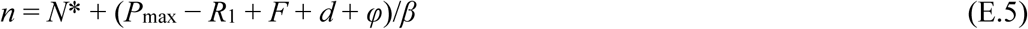

Comparing this with the Nash equilibrium size *n*_Nash_ = *N*\* + (*P*_max_ − *R*_1_)/*β* − 1, expression (E.5) exceeds it by 1 + (*F* + *d* + *φ*)/*β*. A group of strictly interchangeable members therefore reaches the Nash size and stops growing before any member finds it worthwhile to leave alone, and the sub-optimal mode of Case 2 never forms.

This is why the heterogeneity invoked in Case 2 is an assumption of that case rather than a gloss on it. The slopes *α* and *β* are group averages, and the member who dissociates is not the average member but one whose private cost of remaining exceeds the mean by enough to bring (E.5) down to the group’s current size. The rate at which such members arise, together with the residence time of the small groups they found, is what sets the height of the low mode, which is why that height is governed by quantities lying outside the payoff parameters, as stated in Section 3.11.

## Notes

https://doi.org/10.5281/zenodo.20358982

### Competing Interest Statement

The authors have declared no competing interest.

### Summary of Updates

The abstract and introduction have been adjusted to explain that "The models hold each member's standing and resource exploitation strategy constant. While both vary and determine a member's own game and its payoffs, explaining this complexity and its consequences is beyond our scope."

